# Automated SWATH Data Analysis Using Targeted Extraction of Ion Chromatograms

**DOI:** 10.1101/044552

**Authors:** Hannes L. Röst, Ruedi Aebersold, Olga T. Schubert

## Abstract

Targeted mass spectrometry comprises a set of methods able to quantify protein analytes in complex mixtures with high accuracy and sensitivity. These methods, e.g., Selected Reaction Monitoring (SRM) and SWATH MS, use specific mass spectrometric coordinates (assays) for reproducible detection and quantification of proteins. In this protocol, we describe how to analyze in a targeted manner data from a SWATH MS experiment aimed at monitoring thousands of proteins reproducibly over many samples. We present a standard SWATH MS analysis workflow, including manual data analysis for quality control (based on Skyline) as well as automated data analysis with appropriate control of error rates (based on the OpenSWATH workflow). We also discuss considerations to ensure maximal coverage, reproducibility and quantitative accuracy.

## 1. Introduction

Over the past decade, protein analysis strategies based on mass spectrometry (MS) have been steadily gaining popularity. Untargeted shotgun proteomics is currently the most widely used technique for qualitative and quantitative measurements of proteins on a large scale. It allows the detection of hundreds to several thousands of proteins in a single run. However, its focus on high proteome coverage leads to some curtailments in reproducibility, quantitative accuracy, and sample throughput ***(1)***. The targeted proteomic technique selected/multiple reaction monitoring (SRM) alleviates these limitations by focusing the mass spectrometer on a defined set of proteins of interest ***(2)***. SRM excels at consistent and accurate quantification of proteins over large sample sets with coefficients of variation (CV) below 15% and offers the largest dynamic range of all MS-based techniques available today ***(3)***. However, SRM measurements are limited to a few dozens of target proteins per sample injection. Novel MS-based proteomic methods, such as SWATH MS, utilize untargeted, data-independent acquisition (DIA) with targeted data extraction to improve the throughput of targeted proteomics by massively multiplexing the targeted detection of peptides. They thereby combine the comprehensiveness of the shotgun method with the quantitative accuracy and reproducibility of SRM. In terms of quantification reproducibility, SWATH MS performs similarly to SRM and offers a dynamic range of at least three orders of magnitude, positioning it as an ideal technology for large-scale and high-quality proteome measurements.

### SWATH MS

In the SWATH MS workflow, proteins are enzymatically cleaved to produce a mixture of homogeneous peptides and then separated by on-line liquid chromatography (LC) coupled to tandem mass spectrometry (MS/MS). Similar to other DIA methods ***(4***, ***5)***, the mass spectrometer recursively cycles through a large m/z range and co-fragments all peptide ions in relatively large isolation windows (or “swathes”) of several *m/z* (Figure 1A) ***(6)***. For each isolation window, the resulting fragment ions are recorded in a high-resolution composite mass spectrum. Thus, the instrument does not explicitly target single precursors as in shotgun or SRM but rather fragments all precursor ions falling into one isolation window simultaneously. Window size and accumulation time per window are chosen such that the instrument can cycle through all windows relatively quickly, allowing every peptide to be fragmented 8-10 times during its chromatographic elution (Figure 1B). In SWATH acquisition, all ionized species of the sample are thus fragmented and recorded in a systematic, unbiased fashion, independently of their abundance. Compared to shotgun approaches, which record fragment ion spectra based on precursor ion intensity, SWATH acquisition systematically records fragment ion spectra every few seconds, thereby allowing reconstruction of the LC elution profile of each fragment ion (Figure 1B). Similar to SRM, the deterministic acquisition strategy makes SWATH acquisition highly reproducible. The acquisition of full, high-resolution fragment ion data provides additional information compared to SRM, similar to parallel reaction monitoring ***(7***, ***8)***.

**Figure 1.**
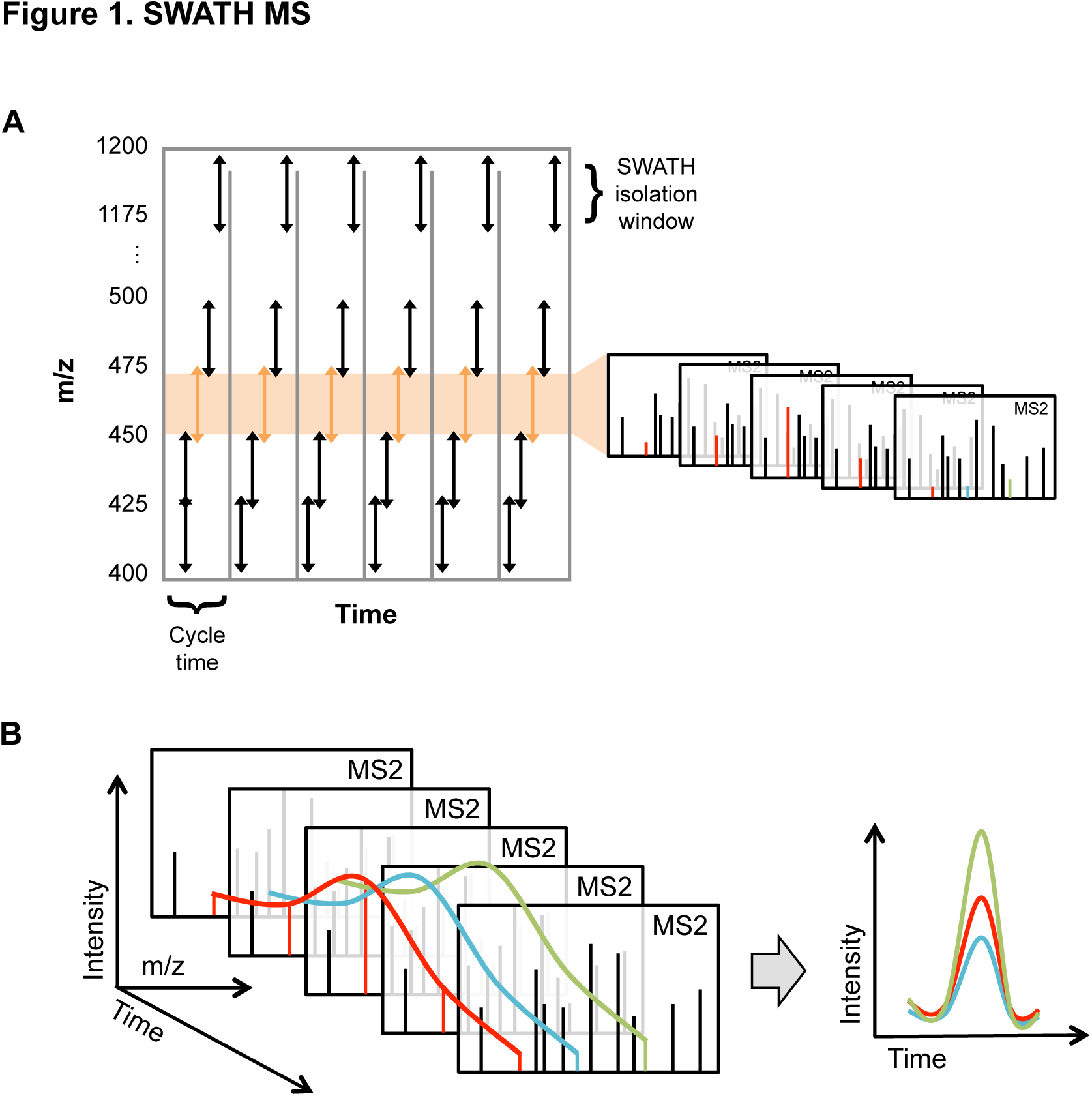
SWATH MS. (A) While peptides elute from the LC, the mass spectrometer cycles through a large m/z range (here from 400-1200 m/z) and co-fragments all peptide ions in relatively large isolation windows (or “swathes”) of several m/z (here 25 m/z) ***(6)***. For each isolation window, the resulting fragment ions are recorded in a high-resolution composite mass spectrum (MS2). For every isolation window, the instrument accumulates fragment ions for 100 ms, for 32 isolation windows covering 400-1200 m/z this results in a cycle time of 3.2 s. At the beginning of each cycle a survey scan (MS1) is recorded (not shown). (B) In SWATH MS, intensities of specific fragment ions are extracted from the highly multiplexed fragment ion spectra in a targeted manner to produce extracted ion chromatogram peak groups.

What distinguishes SWATH MS from other DIA methods is the way the data, i.e. the highly multiplexed fragment ion spectra, are analyzed ***(6***, ***9)***. In SWATH MS, intensities of specific fragment ions are extracted from the highly multiplexed fragment ion spectra in a targeted manner to produce extracted ion chromatograms (XICs) (Figure 1B). These XICs are similar to SRM traces and can be analyzed analogously. This approach facilitates data analysis by reducing the complexity of the data significantly, but requires prior knowledge on the peptides and proteins to be analyzed. This prior knowledge consists of a set of MS coordinates uniquely describing the protein of interest (also called an “assay”). The complete set of coordinates describes (i) which peptides of a protein are most representative, i.e. unique and well detectable, (ii) the elution time of these peptides from the LC, (iii) their pre-dominant charge state, (iv) the most abundant fragment ions formed during fragmentation and (v) their relative intensity. Comprehensive, high-quality assay libraries are a critical prerequisite for the success of a SWATH MS analysis ***(10)***.

### Automated SWATH MS analysis

While small-scale analysis of SWATH MS data can be performed manually (similarly to SRM data analysis), this strategy is not feasible any more for proteome-wide analysis. Automated software packages provide fast, unbiased and comprehensive data analysis solutions using advanced machine learning and data processing algorithms. These software packages are able to automatically import assay libraries, extract chromatographic traces from the raw data and identify peak groups resulting from co-eluting fragment ions (Figure 2). These peak groups are then scored and statistically evaluated using a noise model (negative distribution of scores). This step allows the software to assign each peak group a quality score, which reflects the probability of making a correct identification. Researchers can then use this score to apply an experiment-wide false discovery rate (FDR) cutoff based on how many false quantifications they are willing to tolerate in the final dataset. If multiple runs are present, a cross-run alignment can be performed that integrates all available data to further increase the consistency of the final data matrix (Figure 2).

**Figure 2.**
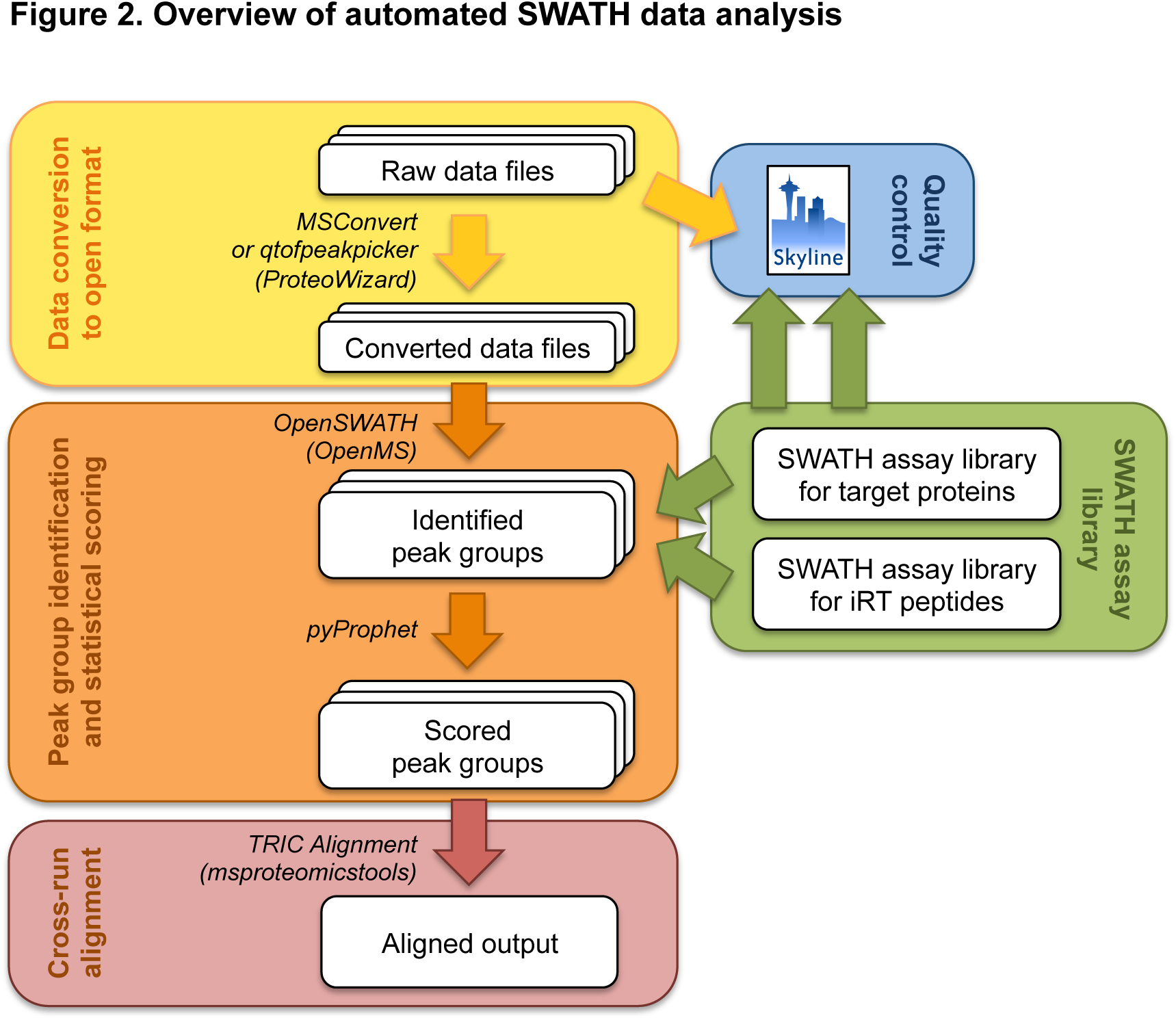
Overview of the workflow for automated SWATH data analysis.

In this chapter we provide a detailed workflow for automated analysis of SWATH MS data using the software suite OpenSWATH, including pyProphet and TRIC Aligner ***(9)***. Because data quality is key for a successful automated SWATH MS analysis, we start with a brief description on how to load SWATH MS data into Skyline ***(11)*** to visually inspect a small number of peptides in every run for quality control.

## 2. Materials

Any files described in this chapter, including example files to test the protocol, can be downloaded from the following website: http://www.peptideatlas.org/PASS/PASS00779. Most of the example files are related to a recent study in *Mycobacterium tuberculosis* that used SWATH MS ***(12)***. The installation of software tools is described for a 64-bit Microsoft Windows 7 system. However, many of the tools can be installed on other platforms, such as Mac OS X and Linux, as well (OpenMS, Python, pyProphet, TRIC aligner). Specific software versions used to develop this protocol are noted in SoftwareVersions.txt. It is important to have enough free disk space available as the data files and the intermediary analysis files tend to be very large (dozens of GB).

1. SWATH data files. A detailed protocol of how to obtain high-quality SWATH MS data on a TripleTOF instrument is provided in Chapter XX by Hunter et al., but other instrument types can be used as well (Note 1). See Note 2 for a discussion of parameters that are important during acquisition, such as LC gradient, SWATH window size, and acquisition time. Importantly, the samples have to contain retention time calibration (iRT) peptides at the time they are measured (Note 3). In this protocol, we assume that the 11 synthetic iRT peptides provided in the iRT-kit by Biognosys have been spiked into each sample ***(13)***. Please note that the AB SCIEX TripleTOF instrument produces two data files per SWATH run, a.wiff and a.wiff.scan file, which should always be stored together (see three.wiff and corresponding.wiff.scan example files).
2. File describing SWATH window setup. The selection of appropriate SWATH windows is an important part of a SWATH MS experiment (Chapter XX by Hunter et al.). For the current protocol, we assume that a fixed window setup (32 windows of 26 m/z each, 1 m/z overlap: 400-425 m/z, 424-450 m/z, 449-475 m/z, … 1149-1175 m/z, 1174-1200 m/z) has been used for data acquisition (see example file SWATHwindows_acquisition.tsv). In contrast to the SWATH windows used for data acquisition, the SWATH windows used for data analysis should not be overlapping. Therefore, the 1 m/z overlap that was used for acquisition among the neighboring SWATH windows is split, such that each window is only 25 m/z (first and last windows are slightly different): 400-424.5 m/z, 424.5-449.5 m/z, 449.5-474.5 m/z, … 1149.5-1174.5 m/z, 1174.5-1200 m/z (see example file SWATHwindows_analysis.tsv).
3. SWATH assay library. In addition to the SWATH data files, the following workflow requires as an input a SWATH assay library, containing pre-computed decoy transition groups as well as assays for the retention time peptides. These assays can be built from experimental data (see ***(10)*** for a detailed protocol) or downloaded from the SWATHAtlas database (see Note 4). As an example library for this protocol we use a comprehensive SWATH assay library of *M. tuberculosis* (see example file Mtb_TubercuList-R27_iRT_UPS_decoy.tsv).
4. iRT peptide SWATH assays. In order to perform targeted extraction, the retention time of each SWATH MS injection needs to be transformed into a normalized retention time space, which is also used by the assay library (Note 3). The protocol we describe here assumes that the 11 synthetic iRT peptides from the iRT-Kit by Biognosys have been spiked into every sample. For this protocol we provide an iRT peptide SWATH assay library in TraML format for OpenSWATH analysis (see example file iRTassays.TraML) and a reduced table for import into Skyline (see example file iRTassays_Skyline.tsv).
5. Skyline. Skyline is a free, open-source software for targeted data analysis of various types of proteomics data ***(11)*** and provides great visualization options. The software can be downloaded from the website: http://skyline.maccosslab.org. It is only available for Microsoft Windows operating systems.
6. Proteowizard. To convert vendor raw data files into an open format, such as mzML or mzXML, we use the msconvert tool, which is part of the Proteowizard software suite (Chapter XX by Mallick et al.). Download Proteowizard from http://proteowizard.sourceforge.net/downloads.shtml and install it on your machine. Due to licensing constraints, the data conversion functionality of ProteoWizard is only available for Microsoft Windows operating systems.
7. OpenMS. OpenMS is a cross-platform and open-source analysis package for mass spectrometric data ***(14***-***17)***. It is specifically suited for automated, large-scale analysis and can be run on any machine, from a Desktop computer to a high-performance Linux cluster ***(18***, ***19)***. Download the appropriate version of OpenMS for your operating system from https://sourceforge.net/projects/open-ms/files/OpenMS/OpenMS-2.0/. Install OpenMS into a local folder, for example C:\Program Files\OpenMS-2.0. On newer Windows versions (Windows Vista, 7 and higher) some of the.NET dependencies will already be installed, which will be indicated during the install and removes the need to download and install them. (The warning during.NET installation “Turn Windows features on or off” can be ignored.)
8. Python. Several scripts used in this protocol require Python 2.7. On Windows, the easiest way to install Python is through Anaconda, which can be downloaded from https://store.continuum.io/cshop/anaconda/. Choose the “Graphical Installer” under the “PYTHON 2.7” tab appropriate for your system (64-bit or 32-bit). Install Anaconda as Administrator on your system (right-click, “Run as” and then select the Administrator user). Select the installation for all users and choose C:\Anaconda2\ as install path (should be the default). Unless you have another Python installation on your system, allow Anaconda to be added to your PATH and register it as the default Python 2.7 installation (these options become available during the installation process).
9. pyProphet. pyProphet is a tool to calculate a discriminating score separating target from decoy assays and to compute a false discovery rate (FDR) cutoff based on this score ***(20)***. Open the “AnacondaPrompt” by opening the Start Menu (Windows icon in lower left corner)and type “Anaconda” in the search field. Right-click the “AnacondaPrompt” entry and select “Run as administrator”. To install the pyProphet package, enter:

~~~
pip install scikit-learn==0.15.2
pip install pyprophet==0.13.3
~~~

If running a 32-bit system, there are no pre-built packages and a working C++ compiler is required on your machine. To get such a compiler, go to http://aka.ms/vcpython27 and download the file called VCForPython27.msi. After installing this file, use the command above to install pyProphet.
10. TRIC feature aligner. The TRansfer of Identification Confidence (TRIC)software tool integrates information from multiple OpenSWATH runs using cross-run alignment and retention time correction. Open the“Anaconda Prompt” by opening the Start Menu (Windows icon in lower left corner) and type “Anaconda” in the search field. Right-click the“Anaconda Prompt” entry and select “Run as administrator”. To install the TRIC aligner package, enter:

~~~
conda install biopython
pip install msproteomicstools==0.3.2
~~~

## 3. Methods

### 3.1 Visualization of selected peptides for quality control

Before proceeding with automated SWATH data analysis, it is crucial to ensure adequate quality of the raw data. Skyline offers great visualization options to assess SWATH data quality and we recommend loading all data files to be subjected to automated analysis by OpenSWATH first into Skyline to manually inspect a few quality control peptides, such as the spiked-in iRT peptides. Optimally, every SWATH data file is loaded into Skyline right after acquisition to monitor LC stability as well as performance of the mass spectrometer.

1. Open a blank Skyline document and go to Settings → Peptide Settings → “Filter” tab. Uncheck the option to “Auto-select all matching peptides” and click “OK”.
2. Go to Settings → Transition Settings → “Filter” tab. Uncheck the option to “Auto-select all matching transitions” and click “OK”.
3. Go to Settings → Transition Settings → “Full-Scan” tab. Set the following parameters: Acquisition method: DIA; Product mass analyzer: TOF; Isolation scheme: From the drop-down menu, select “Add…” and fill the “Edit Isolation Scheme” window as shown in Figure 3A (for this protocol, the window boundaries can be copy-pasted from the file SWATHwindows_analysis.tsv). Resolving power: 15,000 (depends on the state of the instrument during SWATH data acquisition); Retention time filtering: Include all matching scans. Click “OK”.
4. Go to Edit → Insert → Transition list. In the opening window, paste assays of peptides you would like to monitor. For this protocol, the assays for the iRT peptides can be copy-pasted from the file iRTassays_Skyline.tsv. Click “Insert”.
5. Save the Skyline file.
6. Go to File → Import → Results and select “Add single-injection replicates in files” and click “OK”.
7. Select the raw SWATH data .wiff files (the .wiff.scan files are not showing up here, but need to be located in the same folder) and click “Open”. When asked about removing the common prefix of the file name, click “Remove”. Now it will take a while to import all the data depending on file size and number of files.
8. After the data is loaded, arrange the Skyline file such that it is convenient to inspect the runs (Figure 3B). First, get the peak area and retention time overview plots: View → Retention Times → Click “Replicate Comparison” and View → Peak Areas → Click “Replicate Comparison”. Next, arrange the panels by drag-and-drop to the desired location. Then go to “Settings” and click “Integrate all”. Finally, right-click on a chromatogram plot → Auto-zoom X-axis → Best Peak.
9. Inspect the SWATH peak groups to judge performance of LC (peak shape, retention time stability) and mass spectrometer (signal intensity and signal-to-noise ratio for low abundant peaks) over time (Figure 3B). Note that Skyline uses a scoring system to determine the correct peak group but it can still happen that it does not pick the correct one. In these cases, peak boundaries can be changed manually by click-and-drag in the retention time axis of the chromatogram plot. A confidently identified peptide is characterized by a well identifiable peak group near the expected retention time with co-eluting fragment ions that match the relative intensities given in the assay library. More criteria that are typically used to identify correct peak groups are discussed elsewhere ***(9***, ***21)***. For detailed Skyline tutorials please consult the Skyline website (http://skyline.maccosslab.org).

**Figure 3.**
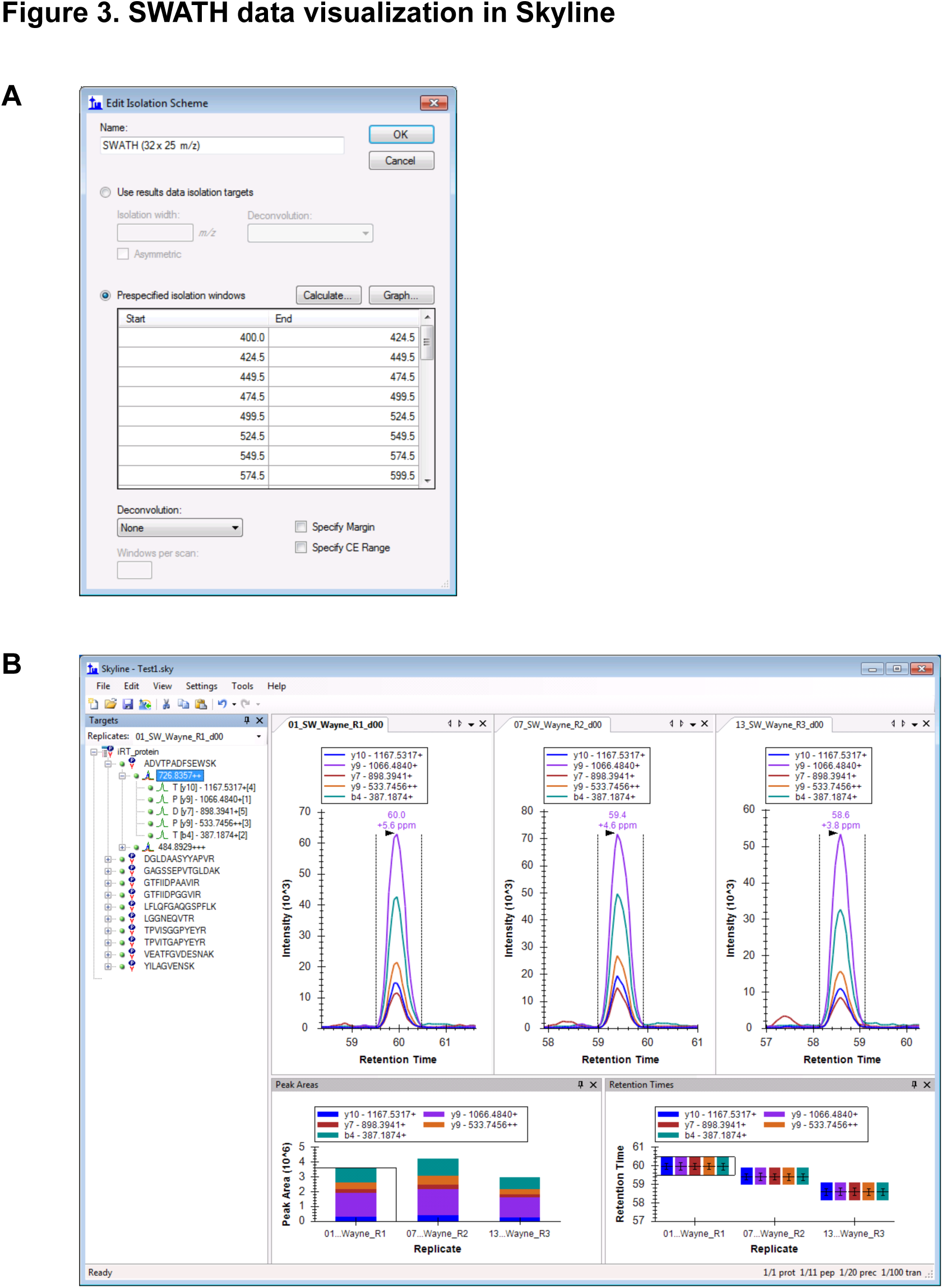
SWATH data visualization in Skyline. (A) Screen shot showing how SWATH windows are defined in Skyline. This is essential for the software to know from which SWATH window the specified fragment ions should be extracted. (B) Example of how to organize the Skyline window for fast and efficient monitoring of large numbers of SWATH runs. The example here shows just three runs, but Skyline can easily handle dozens of runs.

### 3.2 Raw data conversion into mzML

MSConvert, provided through the ProteoWizard software suite (Chapter XX by Mallick et al.), enables conversion of proprietary raw data files (.wiff for AB SCIEX,.raw files for Thermo Fisher instruments) into an open format, such as mzML or mzXML. OpenSWATH can handle both file types, but when running on a Windows PC, the input files need to be in mzML format.

1. Start the software by opening the Start Menu (Windows icon in lower left corner), type “MSConvert” in the search field and click on “MSConvert”.
2. In the MSConvert window, select “List of Files” and use the “Browse” button to locate the raw files to be converted (the.wiff and.wiff.scan files need to be located in the same folder). Click the “Add” button. Choose an output directory using the second “Browse” button. Then set the following parameters: Output format: mzML; Extension: mzML; Binary encoding precision: 64-bit; Write index: yes; Use zlib compression: yes. All other checkboxes should be unchecked and no filters should be set (see Figure 4). While OpenSWATH produces best results on profile data, it is possible to run it on centroided data to reduce disk space and execution time (see Note 5).

**Figure 4.**
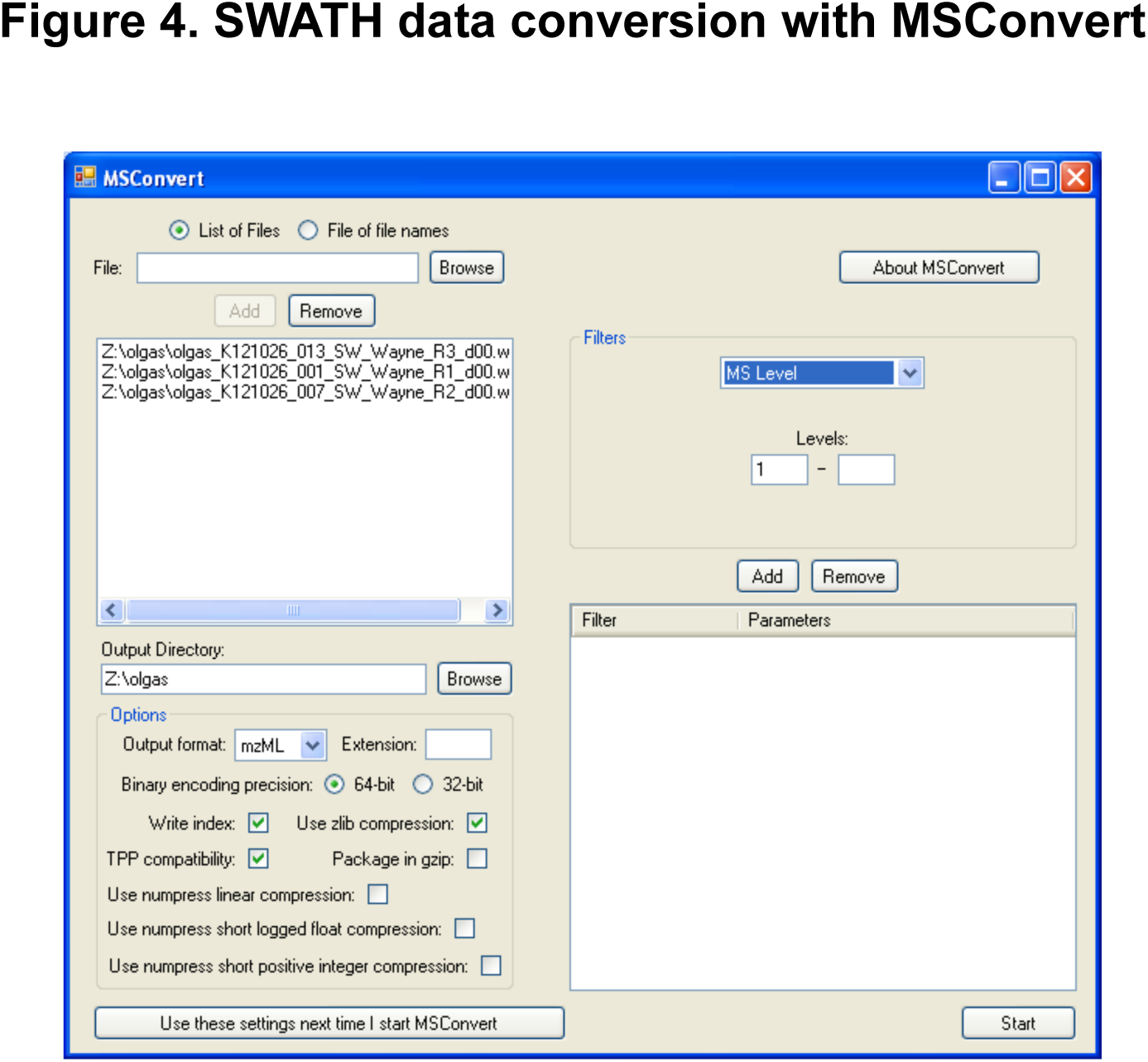
SWATH data conversion with MSConvert. Conversion of raw data to mzML format using the MSConvert GUI, showing the individual options that should be selected during the conversion step for profile-mode conversion.

### 3.3 Automated analysis with OpenSWATH

OpenSWATH can be run as a single command, which executes all steps of the OpenSWATH data analysis pipeline automatically. The individual steps can be controlled through flags passed on the command line. Use the `--helphelp` command to see all options and to learn about particular speed and memory optimizations available. Table 1 summarizes the most important options.

**Table 1:** OpenSWATH parameters

| Parameter | Explanation |
| --- | --- |
| `--helphelp` | To see all options |
| `-out_chrom` | Additionally also output the generated XICs. This option is required if the OpenSWATH results are to be inspected manually with TAPIR **(28)** or for other downstream processing not discussed here. |
| `-swath_windows_file` | Text file specifying the SWATH windows (see example file and main text for more details). |
| `-rt_extraction_window` | The size of the retention time extraction window in seconds (centered around the theoretical retention time calculated using the iRT alignment), usually 300 to 600 seconds are reasonable, but may be adjusted according to the chromatography and the accuracy of expected retention times in the assay library. |
| `-mz_extraction_window` | Size of the extraction window in Dalton. The default value of 0.05 may need to be adjusted depending on the instrument resolution. |
| `-ppm` | Indicates that `-mz_extraction_window` is given in ppm instead of Dalton. |
| `-Scoring:TransitionGroupPicker:min_peak_width` | Minimal peak width in seconds preventing spurious peaks from appearing in the results. Defaults to 14 seconds. |
| `-TransitionGroupPicker:background_subtraction` | Indicates whether background subtraction should be performed during quantification. This option may increase the accuracy of the quantification. |
| `-Scoring:TransitionGroupPicker:PeakPickerMRM:sgolay_frame_length` | Smoothing parameter for Savitzky-Golay smoothing (expressed in number of scans). Increasing this parameter leads to stronger smoothing. Alternatively, Gaussian smoothing may be used by `-Scoring:TransitionGroupPicker:PeakPickerMRM:use_gauss`. |
| `-Scoring:stop_report_after_feature` | Determines the number of peak groups per assay that are reported in the result. A smaller number produces less output; generally 5 to 10 is appropriate. |
| `-batchSize` | Determines the size of the batch of chromatograms that are loaded into memory and analyzed at once. The smaller this number, the less memory is required. |
| `-sort_swath_maps` | This parameter ensures that the SWATH maps in the input are sorted by m/z (this assumes that the swath window file given in `-swath_windows_file` also contains windows sorted by m/z) |
| `-readOptions` | Setting this option to `cache` instructs the software to not load all data into memory at once but to create a local cache of the data. Make sure to specify a directory for the temporary files using the `-tempDirectory` parameter. |
| `-tempDirectory` | Defines the directory to store temporary files (required if `-readOptions cache` is used). |

1. Start the software by opening the Start Menu (Windows icon in lower left corner), type “TOPP” in the search field and click on “TOPP command line” which will start a command prompt pre-loaded with the necessary paths to execute OpenSWATH.
2. OpenSWATH requires four input files: (1) The actual data in mzXML or mzML format (on Windows PC only mzML format is accepted). (2) An assay library containing assays for all target peptides. OpenSWATH can take assay libraries in both tab-separated table (tsv) and TraML format. For more details see Note 4 and Note 6. (3) An assay library containing assays for all iRT peptides (Note 3) in TraML format. (4) A file with a small table specifying the SWATH windows out of which the fragment ion traces should be extracted. Note that the extraction windows are slightly different from the windows specified for data acquisition, i.e. they should not contain overlaps (see example files described in the Materials section). A full command for running OpenSWATH looks like this (enter entire command on a single line):

~~~
OpenSwathWorkflow.exe
-in data.mzML
-tr library.tsv
-sort_swath_maps
-readOptions cache
-tempDirectory C:\Temp
–batchSize 1000
-tr_irt iRT_assays.TraML
-swath_windows_file SWATHwindows_analysis.tsv
-out tsv osw output.tsv
~~~

The output of the OpenSWATH command is a large table containing one scored peak group per row (usually more than one peak group is scored per assay). The properties of each peak group are given in columns (i.e. the assay used to generate the peak group, the retention time, the individual scores etc.).

### 3.4 Error rate estimation with pyProphet

After running OpenSWATH, q-values corresponding to the FDR of peak identification can be estimated with the pyProphet software tool. The only input for pyProphet is the OpenSWATH result table file (osw_output.tsv).

1. Use the same command line window as above and enter (on a singleline):

~~~
C:\Anaconda2\Scripts\pyprophet.exe
--ignore.invalid_score_columns
--d_score.cutoff=0.5>
osw_output.tsv
~~~

This command will output a set of files, of which osw_output_with_dscore_filtered.csv will be used in the next step.
2. pyProphet will also output a table containing the number of true and false positives for different q-value cutoffs (_summary_stat.csv). A q-value cutoff (or FDR) between 1% and 10% is appropriate for most studies. For further information, the pdf report file should be consulted,which contains plots of the fitted distributions as well as ROC curves.This step provides an opportunity to identify common errors, such as using an assay library without decoys or using an assay library unsuited for the measured sample (e.g. from another organism). In asuccessful run, target and decoy distributions should be clearly separated as shown in Figure 5.

**Figure 5.**
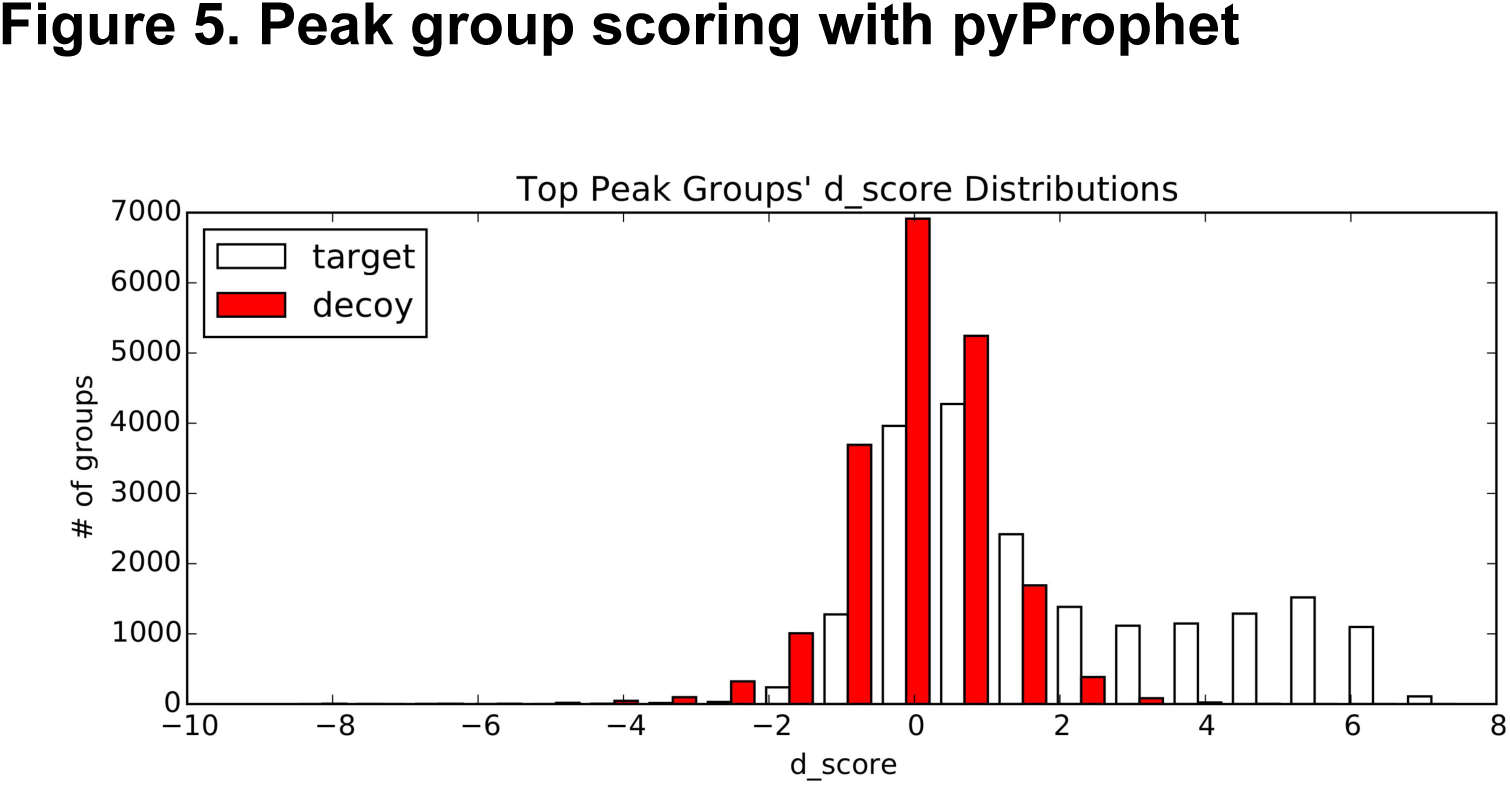
Peak group scoring with pyProphet. To ensure that the FDR estimation step using pyProphet worked properly, the output should be manually inspected to identify potential problems. The plotted distributions should be bimodal and the decoy distribution should be close in shape to the false positive distribution (left part of the bimodal distribution).

### 3.5 Cross-run alignment with TRIC aligner

If multiple SWATH MS runs were analyzed with OpenSWATH, the resulting quantitative data matrix may contain a substantial number of missing values. One way to reduce these missing values is to use the TRIC algorithm to perform cross-run alignment. This algorithm aligns peak groups across all runs using either a reference-based alignment or a tree-based alignment strategy and is implemented in the `feature_alignment.py` tool from the msproteomicstools package. Table 2 describes the individual parameters. For further information you may also consider the online documentation at https://github.com/msproteomicstools/msproteomicstools.

**Table 2:**
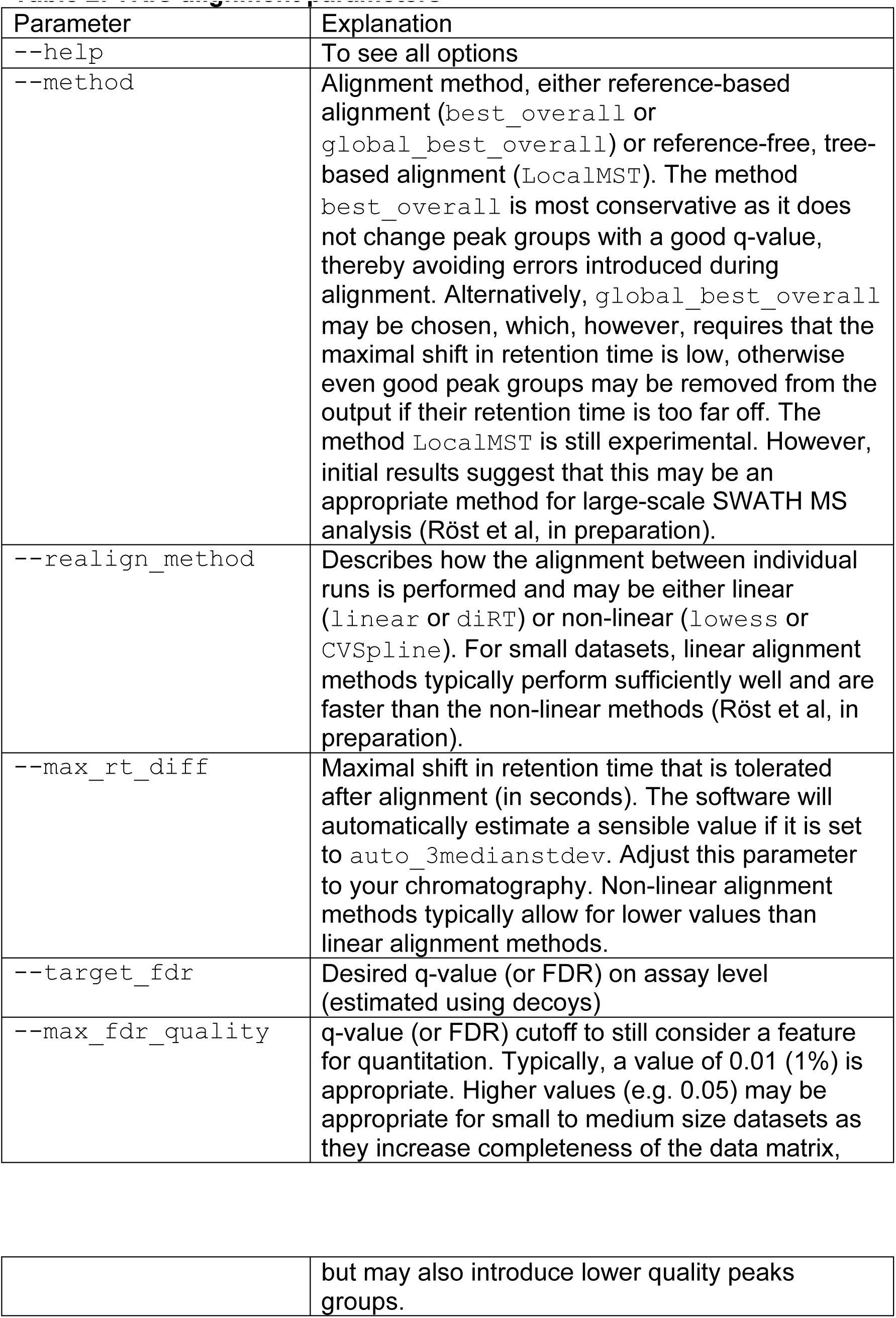
TRIC alignment parameters

1. Use the same command line window as above and enter (on a single line):

~~~
C:\Anaconda2\python.exe
C:\Anaconda2\Scripts\feature_alignment.py
--in file1_with_dscore.csv file2_with_dscore.csv
file3_with_dscore.csv
--out aligned.tsv
--method best_overall
--realign_method diRT
--max_rt_diff 90
--target_fdr 0.01
--max_fdr_quality 0.01
~~~

The command exemplified here will run alignment on three files using linear iRT alignment and pick an appropriate peak group in each run within the aligned window using a reference-based alignment. Only peptides that pass the identification threshold estimated by the TRIC algorithm will be reported in the final output (set to 1% FDR using the `--target_fdr` parameter in the example above). Note that recent results suggest substantial improvements when using reference-free, non-linear retention time alignment and algorithmic improvements may become available in the near future (see https://github.com/msproteomicstools/msproteomicstools, Röst et al., in preparation).

## 4. Notes

1. While SWATH MS was originally developed on an AB SCIEX TripleTOF instrument, SWATH-like data-independent acquisition can also be obtained from other types of instruments. One of the major considerations is the acquisition speed, which needs to allow sufficient sampling during elution of an analyte from the LC column. The software tools described here, i.e. Skyline and OpenSWATH, can both analyze data from multiple vendors, including data acquired on Waters Synapt TOF instruments, Thermo Fisher Q Exactive instruments and AB SCIEX TripleTOF instruments ***(9)***. Multiple groups have reported SWATH-like acquisition on Thermo Q Exactive instruments and successfully used OpenSWATH and other SWATH MS software for data analysis ***(22***-***24)***.
2. SWATH acquisition requires careful optimization of multiple parameters including chromatographic peak width, m/z window width, acquisition time per window and overall m/z coverage. In the initial implementation of SWATH MS, these parameters were set to ca. 30 seconds for the chromatographic peak width, 25 m/z for the window width, 100 ms for the acquisition time per window and a precursor mass range of 400-1200 m/z was chosen, resulting in 32 SWATH windows. Since then, multiple improvements to the scheme have been proposed (Chapter XX by Hunter et al.). It is important to remember that one of the main constraints is to retain sufficient sampling of each peptide precursor during its chromatographic elution. In the original implementation, no more than 100 ms acquisition time could be allowed in order to complete a 32-window cycle within 3.3 seconds. Assuming an average chromatographic peak width of 30 seconds, this would enable the sampling of on average 9 data points across the peak which is sufficient for the OpenSWATH algorithm to reconstruct a peak and perform fragment ion trace cross-correlation. If more powerful chromatographic separation were employed (as offered by UPLC, for example), the other parameters would have to be adjusted accordingly. For example, the acquisition time of each SWATH window could be reduced in order to retain sufficient sampling. Also, the precursor mass range could be decreased or the size of each window increased, both resulting in a smaller number of SWATH windows allowing the instrument to complete each cycle faster. Also, working with flexible window sizes has been shown to improve performance (Chapter XX by Hunter et al.).
3. The presence of iRT retention time reference peptides, so-called iRT peptides, is crucial to calibrate the retention time information present in the SWATH assay library to the retention times recorded in the SWATH data. In principle, any set of well-detectable peptides spanning a wide retention time range can be used ***(13)***. Even endogenous peptides can serve as iRT peptides, making it unnecessary to purchase and spike synthetic peptides into every sample. A set of conserved endogenous peptides found in most eukaryotic samples has recently been described ***(25)***. These Common internal Retention Time standards (CiRT) can be used in the same way as the iRT peptides without the need to spike in additional peptides. However, for some specialized applications, i.e. blood plasma analysis, the CiRT approach may be sub-optimal and the use of the commercial iRT peptides is recommended.
4. Comprehensive, ready-to-use SWATH assay libraries have been published for a number of organisms, including human ***(26)*** and yeast ***(27)***, and can be downloaded from http://www.SWATHAtlas.org. Schubert et al. provide an extensive discussion on various aspects of SWATH assay libraries and provide a detailed protocol on how to build high-quality assay libraries from shotgun data ***(10)***.
5. The OpenSWATH software is designed and tested on profile data and produces optimal results on profile data. However, it is possible to run OpenSWATH on centroided data, reducing the required disk space and execution time of the whole workflow substantially. As centroiding reduces the number of peaks in the data and sometimes may remove low-intensity peaks, the XICs generated by OpenSWATH become less smooth and peak detection becomes less sensitive. If centroiding is performed, it is crucial to compare the results of OpenSWATH to those obtained on profile data on the same dataset to obtain an accurate estimation of how centroiding affects the identification rate. Multiple centroiding algorithms are freely available, including the internal msconvert centroiding (enabled during conversion by the “Filter” settings) as well as the separate executable `qtofpeakpicker.exe` (found in the same folder as `msconvert.exe` after installation of ProteoWizard). The qtofpeakpicker has been specifically designed for data derived from QTOF instruments and we recommend to use it to centroid TripleTOF data ***(10)***. The qtofpeakpicker is run directly on the.wiff files and we recommend using the same parameters as Schubert et al. to generate centroided mzML files ***(10)***: `--resolution=20000 --area=1 --threshold=1 -- smoo thwidth=1.1`.
6. A high-quality SWATH assay library is an essential feature for any targeted analysis of DIA or SWATH MS data. The OpenSWATH software uses this information to extract XICs for the targeted peptide from the appropriate fragment ion spectra and to score the peaks according to how well they match the information in the assay library (expected retention time, fragment ion m/z, etc.). Alongside the assays for the proteins of interest (target assays), the library needs to contain a set of “decoy assays” which represent peptides not present in the sample (usually shuffled target peptide sequences). These assays are scored in the same fashion as the targeted assays and used by pyProphet to estimate the null distribution of different OpenSWATH scores and compute q-values (for FDR control). Having decoys in the assay library is essential for this step to work. Repositories such as http://www.SWATHAtlas.org provide assay libraries that already contain decoys. Please refer to Röst et al. and Schubert et al. for more information on the scoring algorithm and the importance of decoys and FDR control ***(9***, ***10*)**. OpenSWATH accepts assay libraries in both tab-separated table (tsv) and TraML format. For large assay libraries, the tsv format is more memory efficient than the TraML format. OpenMS comes with the tools `ConvertTraMLToTSV` and `ConvertTSVToTraML`, which enable conversion between the two formats (regardless of the names of these conversion tools, the file ending for the table format has to be.csv, while the actual data format is tab-separated).

